# A combined characterization of coelomic fluid cell types in the spiny starfish *Marthasterias glacialis* – inputs from flow cytometry and imaging

**DOI:** 10.1101/2020.06.20.163196

**Authors:** Bárbara Oliveira, Silvia Guatelli, Pedro Martinez, Beatriz Simões, Claúdia Bispo, Claúdia Andrade, Cinzia Ferrario, Francesco Bonasoro, José Rino, Michela Sugni, Rui Gardner, Rita Zilhão, Ana Varela Coelho

## Abstract

Coelomocytes is a generic name for a collection of cellular morphotypes, present in many coelomate animals, that has been reported as highly variable across echinoderm classes. The roles attributed to the major types of the free circulating cells present in the coelomic fluid of echinoderms include immune response, phagocytic digestion and clotting. The main aim of the present study is the thorough characterization of coelomocytes present in the coelomic fluid of *Marthasterias glacialis* (class Asteroidea) through the combined use of flow cytometry (FC) and fluorescence plus transmission electron microscopy. Two coelomocyte populations (here named P1 and P2) were identified by flow cytometry and subsequently studied in terms of abundance, morphology, ultrastructure, cell viability and cell cycle profiles. Ultrastructurally, P2 diploid cells showed two main morphotypes, similar to phagocytes and vertebrate thrombocytes, whereas the small P1 haploid cellular population was characterized by a low mitotic activity, relatively undifferentiated cytotype and a high nucleus/cytoplasm ratio. These last cells resemble stem-cell types present in other animals. P1 and P2 cells differ also in cell viability and cell cycle profiles. Additionally, two other morphotypes were only detected by fluorescence microscopy and a third one when using a combined microscopy/FC approach.

## Introduction

*Marthasterias glacialis* (Linnaeus, 1758) is a member of the class Asteroidea (phylum Echinodermata), commonly known as starfish. Asteroids are known for their remarkable regenerative abilities (1), (2) which allow the animal to restore and regrow lost body parts after injury, providing an ecological advantage in environments shared with predators. The regenerative process involves the mobilization of many cell types to the area of injury, cells that have to perform two major activities: monitor and clear environmental pathogens that suddenly have access to internal fluids, plus the healing of the wound and the subsequent rebuilding of missing structures. It has been known for some time that coelomocytes, the circulating cells present in the coelomic cavities, have a critical role in all these processes (3). Coelomocytes are mostly known as effectors of the immune response and, thus, they are key contributors to the monitoring and restoration of missing tissues (4). How these processes occur is still a matter of debate, due to the still limited knowledge of the fluid’s composition and the mechanistic/molecular basis of coelomocytes functions.

In echinoderms the coelomic fluid completely fills the coelomic spaces of their body, including the perivisceral coelomic cavities, the water vascular system, and the perihaemal systems (5); (6); (7). Coelomocytes perform diverse immune functions, such as formation of cellular clots, phagocytosis, encapsulation and the clearance of bacteria and other foreign materials (8); (7). After starfish traumatic injury or autotomy, coelomocytes rapidly aggregate in the damaged area, forming a clot that seals the inner environment from the external milieu, thus contributing to avoid the loss of coelomic fluid (haemostasis) (8); (9).

Despite what has been reported in several studies, the tissue where coelomocytes originate is still mostly unknown. Different authors have proposed that these cells originate in the axial organ (10), the Tiedemann’s bodies (11) or the perivisceral coelomic epithelium (CE) (12); (13). The latter hypothesis is currently considered the most plausible and is supported by several experimental results that show the release of free circulating cells from the CE in response to injury or to the introduction of foreign particles (14); (12).

While the activities of main coelomocytes populations are known (mostly in immune response, as mentioned above), the number, types and specific contribution of each morphotype to the physiology of the animal in any single species are still poorly understood. Indeed, and due to several reasons like the diverse descriptions of their morphology given by many authors, the use of different names for similar cellular phenotypes, the differences arising from alternative coelomocyte’s handling protocols and the diversity of methodologies used for their characterization, these circulating cells have been subjected to various classification schemes. The main consequence of using these different approaches has been an intrinsic difficulty in homologizing cell types within and across different echinoderm classes. For example, Parrinello (1995)(8) identified in all echinoderms six main categories (amoebocytes, spherule cells, vibratile cells, haemocytes, crystal cells and the progenitor lymphocyte-like cells), whereas Smith and collaborators (2010) (7) identified in sea urchins only three categories of coelomocytes (phagocytes, spherule cells or amoebocytes, and vibratile cells). In the starfish *Asterias rubens*, Gorshkov and collaborators (2009) (15) detected two types of cells that were named just mature and immature coelomocytes. In our model starfish *M. glacialis*, Franco (2011) (16), using light microscopy, reported four types of coelomocytes: spherule cells, vibratile cells, amoebocytes and phagocytes (the latter showing petaloid or filopodial alternate morphologies). Giving this range and diversity of morphologies, it is nowadays unanimously accepted that the types and characteristics of coelomocytes vary among the five echinoderms classes (3). Strikingly, as today, a shared agreement in the classification of the intra-class (*i*.*e*. Asteroidea) coelomocytes has not yet been achieved.

Remarkably, different research teams have mostly focused on the classes of Holothuroidea and Echinoidea for the study of coelomocytes and their immune function, with the other classes remaining, in comparison poorly known. In order to contribute to a deeper understanding of the cellular components present in the coelomic fluid of an asteroid, we carried out a comprehensive analysis and the characterization of coelomocytes of the starfish *M. glacialis*. These results are here discussed with the main aim of clarifying what is the diversity of coelomocytes present in asteroidean. Whenever the current knowledge allows, we take in our discussions an inter-class perspective.

## M&M

### Animal collection and maintenance

Adult specimens of both sexes of the starfish *M. glacialis* were collected at low tide on the west coast of Portugal (Estoril, Cascais). The animals were transferred to the “Vasco da Gama” Aquarium (Dafundo, Oeiras) where they were kept in open-circuit tanks with re-circulating seawater at a temperature of 15°C and a salinity of 33‰. They were fed *ad libitum* with a diet of mussels collected in the same site. All specimens were maintained in the same conditions throughout the whole experimental period to avoid variability due to abiotic factors (e.g. salinity and temperature).

### Coelomic fluid collection

To minimize contamination, coelomic fluid was collected with a 21-gauge-butterfly needle, and transferred directly to a Falcon tube kept on ice. The needle was inserted in the tip of the starfish arm to avoid the disruption of internal organs (*i*.*e*. pyloric caeca and gonads) while coelomic fluid was collected. Coelomic fluid samples were transported in ice from the ‘Vasco da Gama’ Aquarium to the laboratory for all downstream experiments. To minimize coelomocytes’ aggregation, all samples were carefully re-suspended with a micropipette.

### Flow cytometry (FC)

A preliminary filtration step, through a 40 μm mesh, was performed before FC analysis to avoid capillary clotting. Samples were run through a CyAn ADP Flow Cytometer Analyser (Beckman Coulter) and data were analysed using a Summit™ software package. Cells were sorted in FACSAria I (BD Biosciences), a 3 laser High Speed Cell Sorter with BD FACSDiva™ (BD Biosciences) software. Lasers and filters λ, respectively, for the analyser and sorter flow cytometers were for DRAQ5 642; 665/20 nm and 633; 660/20 nm, and for DAPI 405; 450/50 nm and 407; 450/40 nm. Data were further analysed using a FlowLogic Software (Inivai Technologies).

### Imaging flow cytometry – staining and image acquisition

Two hundred μL of coelomic fluid (average number of cells: 90,000) were filtered just before each imaging flow cytometry assay to remove cellular aggregates. Vacuolar membranes and nucleus were stained, respectively, with 0.5 μL of FM4-64 (Molecular Probes, #T3166) at a concentration of 5 μg/mL and 3.0 μL of DAPI (Molecular Probes, #T1316) at a concentration of 5 mg/mL.

Coelomocyte images were acquired using the INSPIRE software of the ImageStreamX Mark II imaging flow cytometer (Amnis Corporation, Seattle, WA) at the Instituto de Medicina Molecular João Lobo Antunes, Faculdade de Medicina, University of Lisbon, Portugal. Cells were imaged with 60x magnification, using a 375 nm laser at 40 mW for DAPI excitation with a detection window of 435-505 nm (channel 7) and a 488 nm laser at 15 mW for FM4-64 excitation with a detection window of 642-745 nm (channel 5). Brightfield images were acquired in channels 1 and 9. Imaging flow cytometry data was analysed with the IDEAS (v6.2) software, following the gating strategy described in the Results section.

### Cell viability assay

To evaluate cell viability, fresh cells were stained with DAPI (Molecular Probes) and DRAQ5 (Molecular Probes) and analysed using a CyAn ADP Analyser. To check for consistency other DNA dyes for dead/permeabilized cells were tested: propidium iodide (PI; Thermo Fisher), DRAQ7 (Thermo Fisher) and 7-ADD (Thermo Fisher). Final concentrations for used dies were: 0.5μg/mL, 0.05μM, 1μg/mL, 0.03μM and 0.5μg/mL, respectively.

### Cell cycle analysis

To study cell cycle progression, an initial cell permeabilization step is needed. The protocol used was adapted from (17). Cells from the coelomic fluid were collected by centrifugation at 1000 xg during 5 min, at 4°C. Pelleted cells were washed once in 3.5% (w/v) Artificial Sea Water (ASW; prepared from Sea Salts, Sigma). Cold 70% (v/v) ethanol was added, dropwise, to the pellet while slowly vortexing. After at the latest 24 hours, cells were rinsed twice in ASW and re-suspended in a solution: 5% RNAse at 100 μg/mL in 85% ASW and 10% PI at 1 mg/mL. Samples were kept protected from light and transferred to a CyAn ADP flow cytometer (laser λ set at 488 nm and filter set at λ 613/620 nm). Data analysis was carried out with the Dean Jett Fox algorithm of the FlowJo software.

### Fluorescence microscopy

In order to analyse the morphology of the cells present in the coelomic fluid, 199 μL were taken off from the cell suspension pellet. A staining protocol with DAPI and FM4-64 (Molecular Probes), as that used for FC-imaging, was adjusted. One μL of FM 4-64 was added directly to the coelomic fluid and incubated for 2 min. Ten μL of this suspension were deposited per well of a microscope slide (Marienfeld Superior), 1 μL of DAPI was added and the mixture homogenized. Microscopy images were acquired by a LEICA DM 6000B with an amplification of 100×1.6, using the software Metamorph, version 7.61.0 (copyright 1992-2009 MDS AT) and processed with the same software.

### Ultrastructural analyses

Coelomic fluid was collected with a needle as previously described and directly dropped into the fixative solution (2% glutaraldehyde and 1.2% NaCl in 0.1 M sodium cacodylate buffer at 4°C) where it was left for at least 2 h. Throughout the protocol changes of solutions were carried out by centrifugation of cells at 400-600 xg for 5 min and subsequent re-suspension of the pellet in the new solution. Cells were washed few times in 0.1 M sodium cacodylate buffer and left in the same buffer overnight at 4°C, post-fixed with 1% osmium tetroxide in 0.1 M sodium cacodylate buffer at room temperature (RT) for 2 h and washed several times with distilled water. They were then left in 2% uranyl acetate in 25% ethanol for 2 h at RT and subsequently dehydrated in an increasing series of ethanol solutions (25%, 50%, 70%, 90%, 95%, 100%), washed in propylene oxide and in a gradual mix of epoxy resin (Epon 812-Araldite) and propylene oxide in different proportions (1:3, 1:1, 3:1). These cells were eventually embedded in fresh resin (Epon 812-Araldite).

Ultrathin sections (between 70 and 90 nm in thickness) were cut with a glass or a diamond knife from cellular pellets using a Reichert-Jung Ultracut E microtome. They were mounted on copper grids (150 and 400 mesh) and stained with a solution of 1% uranyl acetate in distilled water and then with a lead citrate solution. Ultrathin sections were observed and photographed using a CM10 Philips transmission electron microscope (TEM) equipped with a Morada Soft Imaging System digital camera operated with an Item Software.

## Results and Discussion

### Coelomocyte fractionation using flow cytometry

The cytometry analysis of the cells isolated from *M. glacialis* coelomic fluid reveals two clearly distinguishable cell populations (here named P1 and P2) (Figure 1A). These two populations differ in the value of their light scattering, where P2 cells show higher values than P1 cells, probably a direct consequence of having a more complex/structured surface (including particulate material such as inclusions or granules) and richer internal cyto-architecture (Figure 1B). P2 population represents 50-80% of total coelomocytes. A maximum of 20% of these cells are osmotically compromised, displaying lower DRAQ5 incorporation/fluorescence values – P2low in Figure 1A. In a previous study with FC done in the asteroid *Leptasterias polaris* only a single population of coelomocytes was observed (18). In specimens of *Asterias rubens* injected with artificial sea water, a main cluster, attributed to amoebocytes, was originally described, which was further sub-divided in three cellular groups that were differentially affected by bacterial challenge (19). FC profiles similar to ours were found by Xing and collaborators (2008) (20) for the circulating cells of the sea cucumber *Apostichopus japonicus* (Holothuroidea) and by Romero and collaborators (2016) (21) in the sea urchin *Paracentrotus lividus* (Echinoidea). These simple cytometric patterns are consistent with several reports (11), (19) that detect phagocytes as the most abundant, or even the only circulating coelomocyte type in asteroids. A higher cellular diversity was described later by Smith and collaborators (2019) (22) in *Strongylocentrotus purpuratus*. In that manuscript, and using FC, live cells were selected based on exclusion of propidium iodide. The authors were able to discriminate among five populations of coelomocytes (based on their side and forward scattering patterns). Phagocytes (or phagocyte-like) represent in this case a heterogeneous class contributing up to 70% of the circulating cells.

**Figure 1.**
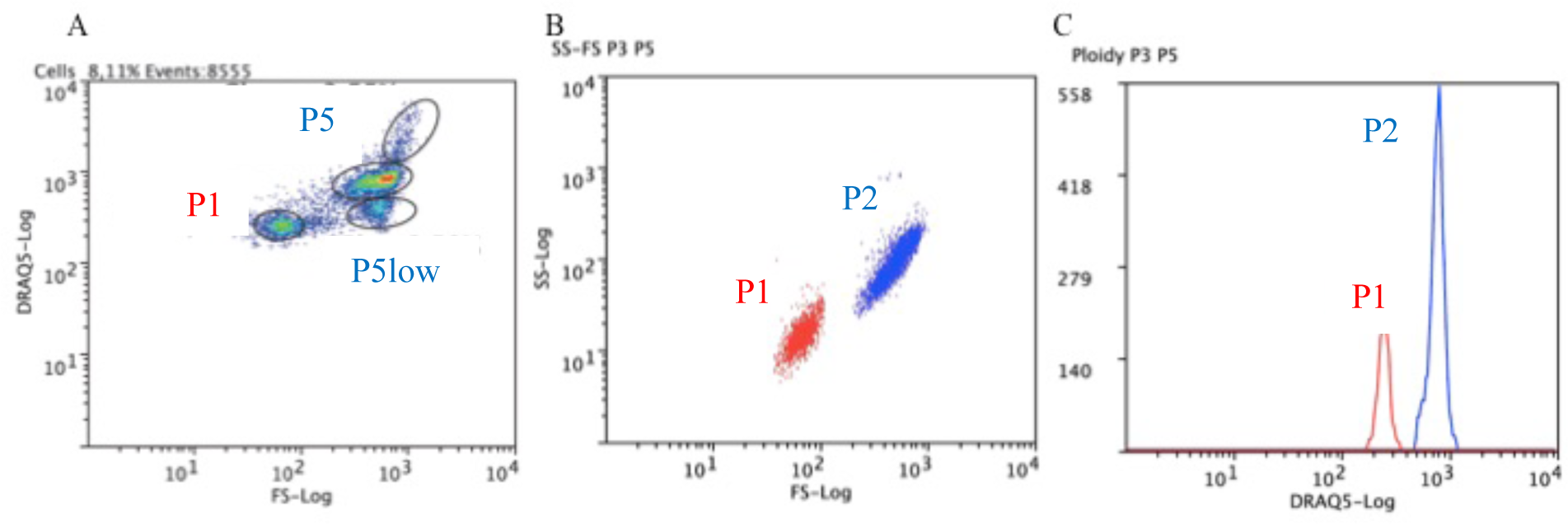
Flow cytometric analysis of circulating coelomocytes. Coelomocytes were stained with DRAQ5. **A**. Representative dot plot of coelomocytes with cell populations gated. **B**. Overlap of dot plots was created to show representative forward and side scatter properties of P1 (red) and P2 (blue) cell populations. **C**. Overlap of P1 (red) and P2 (blue) cell populations histograms was created to show cell populations ploidy.

So far, for 13 starfish species belonging to four families, it has been reported that the genome is packed into 44 diploid chromosomes (23). We determined the values of ploidy for our two identified populations and have found that P2 cells have, approximately, twice more DNA content than P1 cells (Figure 1C). A similar study carried out by Fafandel and collaborators (2008) (24) reported also the existence of two populations with diverse ploidy, in which the DNA content ratio between them was around 4. The authors suggested that some cells in the high-DNA population had probably engulfed other, unknown, nuclei. Although this type of situation was only reported during cell apoptosis (25), this result fit with the presence in the samples of haploid and diploid cells, as we suggest for P1 and P2 cells.

Direct comparison of echinoid coelomocyte types identified by FC with those of Asteroidea is difficult given the known variability of coelomocyte types among echinoderm classes (7); (22) and by the diversity of protocols, for instance, the specific lectins-labelling method used by Liao and collaborators (2017) (26).

Due to their clotting functions, coelomocytes tend to aggregate in the collection process. As a consequence of this we cannot exclude the possibility that a fraction of coelomic cells were retained during the filtration through a 40 μm nylon cell strainer step we had to perform in advance to the flow cytometry analysis. Moreover, the inclusion of anticoagulant solutions such as 1.9% sodium citrate (1:1) (27); (28) and isotonic 1:1 anticoagulant buffer to the suspension (0.5 M NaCl, 5 mM MgCl_2_, 20 mM HEPES and 1 mM EGTA pH=7.5) (29) had the unwanted effect of producing an extensive loss of cells and of compromising of the morphological integrity of the remaining; mainly those of the P1 cell population, as checked by using FC (Figure S1). Unexpectedly, a side effect of using the isotonic anticoagulant buffer was an increase in the number of cell aggregates generated in the sample, as we detected them by FC (Figure S1). In order to minimize these effects of aggregation, the careful coelomocyte resuspension and subsequent filtering of cell suspension were implemented, with the aim of avoiding the use of any anticoagulant in the final protocol. This decision follows the recommendation of Matranga and collaborators (2005) (30) that stated: “to suggest a nomenclature that reflects the actual morphology of the cells requires the immediate observation of fresh and alive cells, just taken from the sea urchin without any addition of anti-coagulant solutions”.

### Evaluation of cell viability

Samples of collected coelomic fluid also carry cellular debris, which interfere with the visualization of coelomocyte populations in cytometry. The use of DRAQ5, a lipophilic and membrane-permeable DNA dye, that has been used for the detection of live cells (31), allowed, up front, to discriminate coelomocytes from cell debris. Additionally, and in order to better evaluate coelomocyte population viability, we introduced the use of compatible exclusion dyes (*i*.*e*. reagents that are excluded by healthy cells owing to the integrity of their plasma membrane and that also do not interfere with DRAQ5 fluorescence or compete for its DNA binding activity). DAPI co-staining fulfilled these criteria revealing that the P1 cell population incorporated it readily, whereas the P2 cell population seemed more refractory and thus, represented a population with relatively few DAPI positive cells (Figure 2). A possible interpretation of the labelling patterns is that, on average, P1 cells exhibit a relative low degree of viability, whereas the majority of P2 cells are viable. Although the low P1 viability does not change much when kept outside the organism over a period of one day, the viability of P2 seems to decrease sharply during the same period. To test if cellular permeability to DAPI is only due to a loss of viability (32), alternative exclusion dyes (PI, DRAQ7 and 7-ADD) were tested. In all cases, the results were almost identical to those obtained with DAPI (data not shown).

**Figure 2.**
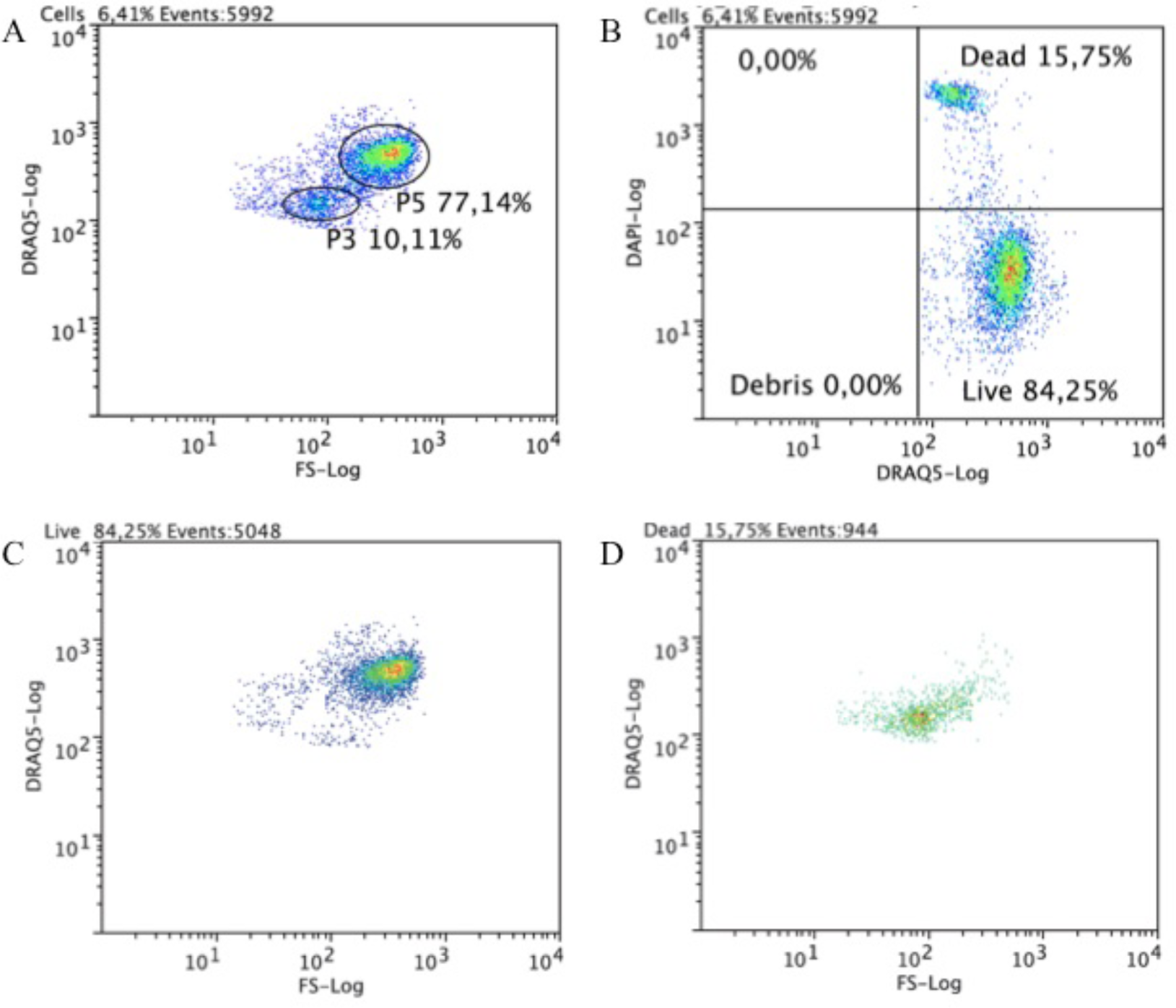
Flow cytometric analysis of circulating coelomocytes viability. Cells were stained with DRAQ5 and DAPI. **A**. Representative dot plot with P1 and P2 cell populations gated. **B**. A quadrant plot of coelomocytes showing DAPI+ (dead) and DRAQ5+ DAPI-(live) coelomocytes. **C**. A dot plot showing that in live cells quadrant, these belong mainly to P2 cell population. **D**. A dot plot showing that in dead cells quadrant they belong mainly to P1 cell population.

After sorting, preliminary fluorescence microscopy tests were carried out to characterize the cellular morphologies of the two detected cell populations. P1 cells were less abundant than P2 and showed an average size of 4 μm, with spherical morphology. Instead, P2 cells presented a diversity of morphotypes with diameters that ranged from 3 to 10 folds those determined for the P1 cells. This P2 diversity of morphologies could also be associated to the particularities of the sorting process. In fact, by higher magnification fluorescence microscopy, using DAPI and FM4-64 to specifically label DNA and cell membranes respectively, it was possible to confirm that most sorted cells were lysed with the total number of intact ones being remarkably low. These effects were also observed when cells were centrifuged and fixed 4 h after collection. The impact of experimental manipulations was not homogenous across the different cell types. Based on this observation no further cell sorting was performed before microscopy assays ensuring that a morphological characterization could be performed without much distortion. Moreover, and to avoid cells’ manipulation, labelling dyes were added directly to the extracted coelomic fluid.

### Analysis of the cell cycle

To study the cell cycle profile, and to determine the fraction of cells going through the different phases, cells were loaded in the cytometer after fixation plus an extensive DNA labelling period with PI to ensure proportionality. FC profile obtained for PI labelled cells mimic that observed with DRAQ5 staining. Data were acquired in linear mode, in order to detect n-fold differences in DNA content. We detected that the P2 fraction of cells was predominantly in the first phase of cell cycle – G0G1 with a 2n DNA content (Figure 3A). P2 cells also seemed to be going through a regular cell cycle, with fractions of cells distributed in all phases. On the other hand, P1 cells are haploid and were all, seemingly, in a unique phase of the cycle (Figure 3B). The simple/single profile of the P1 cells can be ascribed to either a quiescent state (known as G0 phase) or to a terminally differentiated one. In contrast to P2, P1 cells do not seem to be going through division.

**Figure 3.**
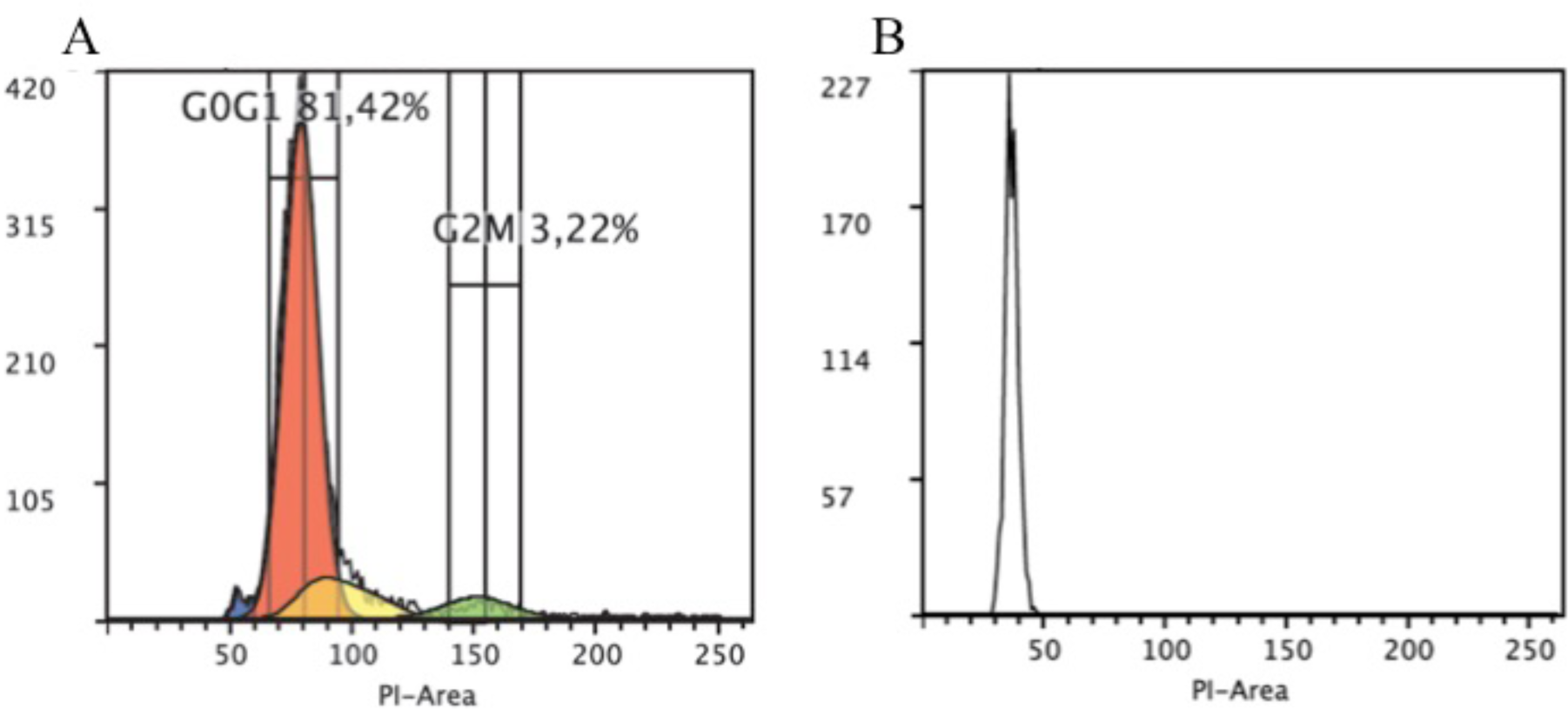
Flow cytometric cell cycle histograms. Cells were fixed with 70% ethanol and stained with PI **A**. Representative histogram of P2 singlets cell cycle showing different phases: sub-G0/G1 (blue) G0/G1 (red), S (yellow) and G2M (green). **B**. Representative histogram of P1 cell population. PI fluorescence is proportional to DNA content.

Interestingly, Fafandel and collaborators (24) reported that in the red starfish *Echinaster sepositus* circulating coelomocytes do not go through mitotic divisions, suggesting that all coelomocytes originate from the coelomic epithelium and are, thus, post-mitotic once they enter in the fluid. Their results contradict ours, although it should be taken into account that the DNA intercalator used in these previous studies (DAPI) stains only (or mostly) dead cells; the reason why in our assays we decided to use PI instead, which binds DNA more extensively, either in dead or alive cells.

### Coupling FC coelomocyte populations fractionation and morphologic characterization by Imaging FC

Imaging flow cytometry assays were performed using stained coelomocyte suspensions prepared by adding 0.5 μL and 3 μL of FM 4-64 and DAPI, respectively, to 200 μL of a filtered coelomocyte suspension sample.

Only focused cell images were selected for further analysis, using the Gradient RMS function in IDEAS to distinguish between focused (high gradient RMS value) and unfocused images (low gradient RMS value). A histogram plot of DAPI staining reveals the same two populations found with DRAQ5 labelling in traditional FC: a P1 population with lower DNA content and a P2 population with higher DNA content (Figure 4A). Whereas P1 corresponds to smaller cells (average cell area = 72 ± 21 μm^2^) with lower FM 4-64 intensity, P2 cells corresponds to bigger cells (average cell area = 226 ± 78 μm^2^) with higher FM 4-64 intensity (Figure 4B). Notice that P2 cells are also more heterogeneous, displaying higher variation in cell size (minimum area = 50 μm^2^; maximum area: 443 μm^2^). In opposition to what was described in *Asterias forbesi*, where only phagocytes were identified (3), several cell morphologies and sizes were distinguished in *M. glacialis* coelomic fluid. The single cell images obtained by imaging flow cytometry show that P1 cells are mostly round, with varying nuclei size (Figure 4C), whereas P2 cells correspond to a mix of different cell morphologies: regular, petaloid, filopodial and big granulated cells (Figure 4D).

**Fig 4.**
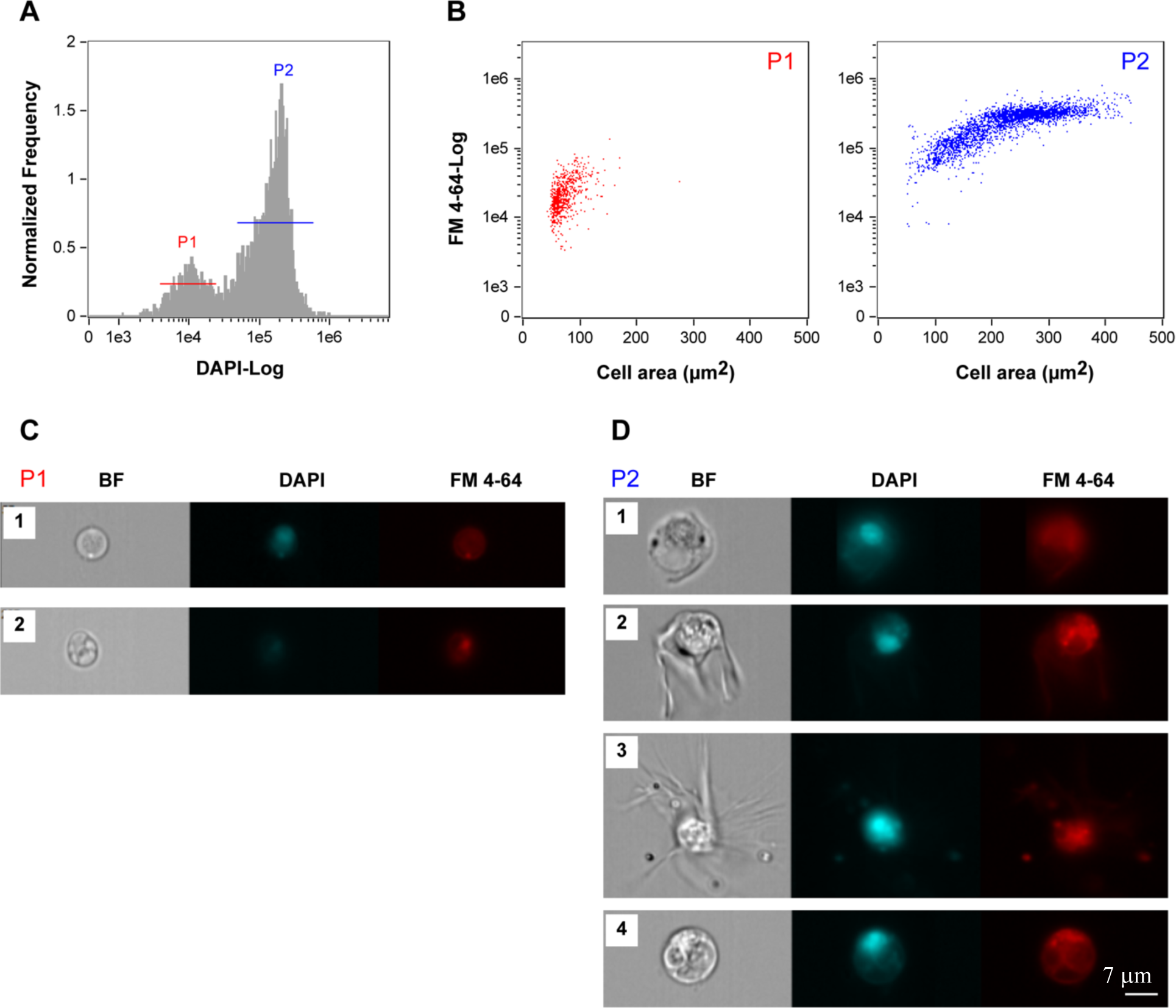
Imaging flow cytometry analysis of circulating coelomocytes. Coelomocytes nucleus and membranes were stained, respectively, with DAPI and FM 4-64. Representatives histogram (**A)** and dot plots (**B)** for P1 and P2 singlets of gated coelomocyte populations, excluding dead cells. Bright field and fluorescence microscopy images for the several morphotypes detected within each coelomocytes populations, P1 (**C**) and P2 (**D**). Coelomocytes nucleus and membranes were stained, respectively, with DAPI (blue) and FM 4-64 (red). C1. P1 cells displaying a nucleus occupying the majority of the cell space; C2. P1 cells displaying a smaller nucleus to cell ratio; D1. P2 regular; D2. P2 petaloid and D3. P2 filopodial coelomocytes; D4. Big granulated cell.

These cell morphologies correspond to most of the cell types observed with high resolution fluorescence microscopy (Figure 5) and electron microscopy (Figure 6), as described below in this manuscript. Combined imaging FC/microscopy results allowed overcoming the difficulties with morphology characterization of FC populations due to coelomocyte susceptibility to sorting. This methodology provided us with extensive cellular information that can be used, eventually, to follow changes within these populations as triggered by a diversity of challenges.

**Fig 5.**
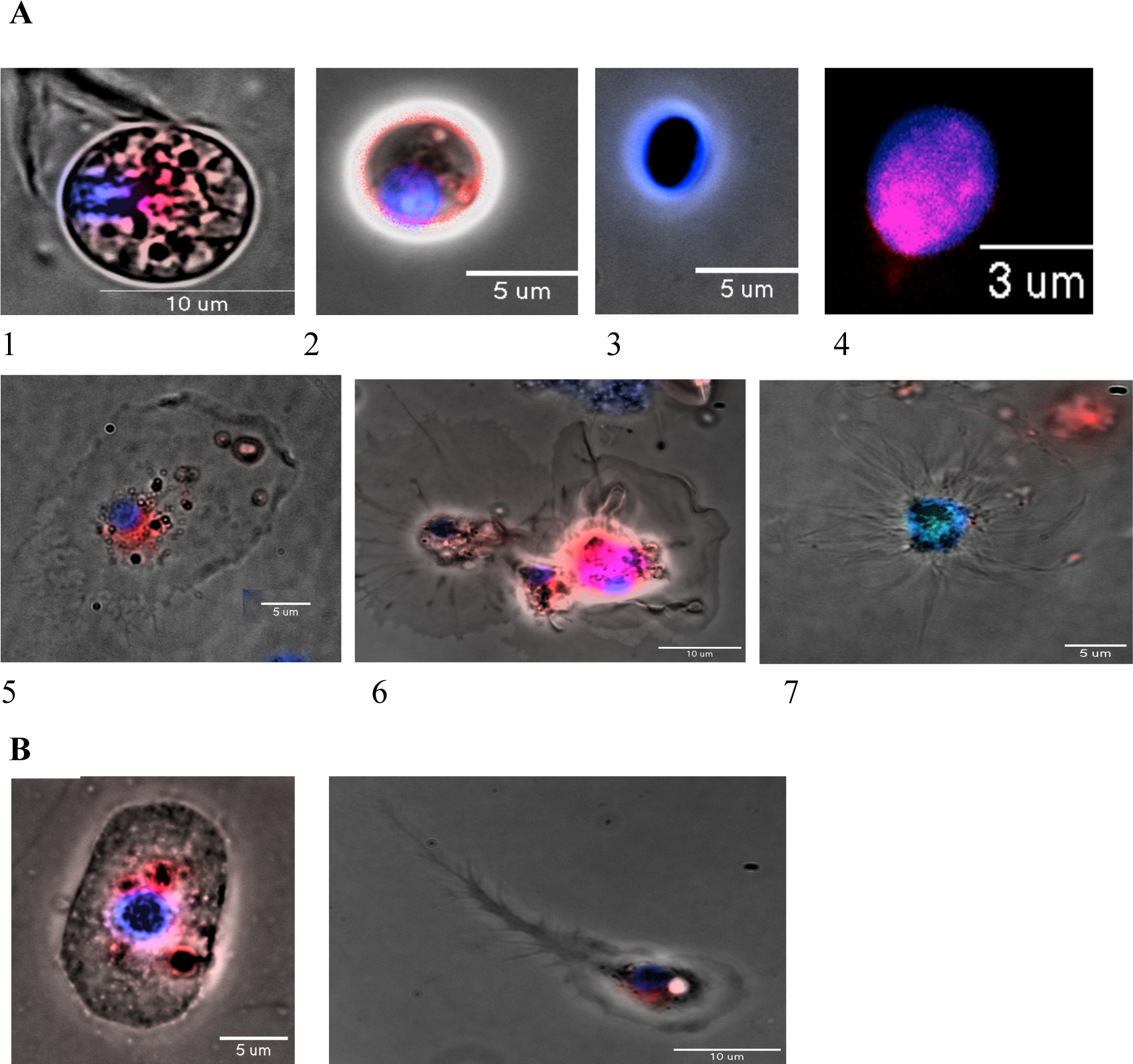
Characterization of coelomocytes morphologies by fluorescence microscopy. **A)** Two morphotypes were identified within P1 cell population **1**,**2**. with a smaller nucleus to cytoplasmic ratio and **3**,**4**. with a nucleus occupying the majority of the cell space. Three morphotypes were observed within P2 cell population **5**. Regular, **6**. Petaloid, and **7**. Filopodial. **B. 1, 2** Representatives of novel P2 cell morphologies Specific staining for DNA and cell membranes was performed with DAPI (blue) and FM4-64 (red) fluorescent dyes.

**Fig. 6.**
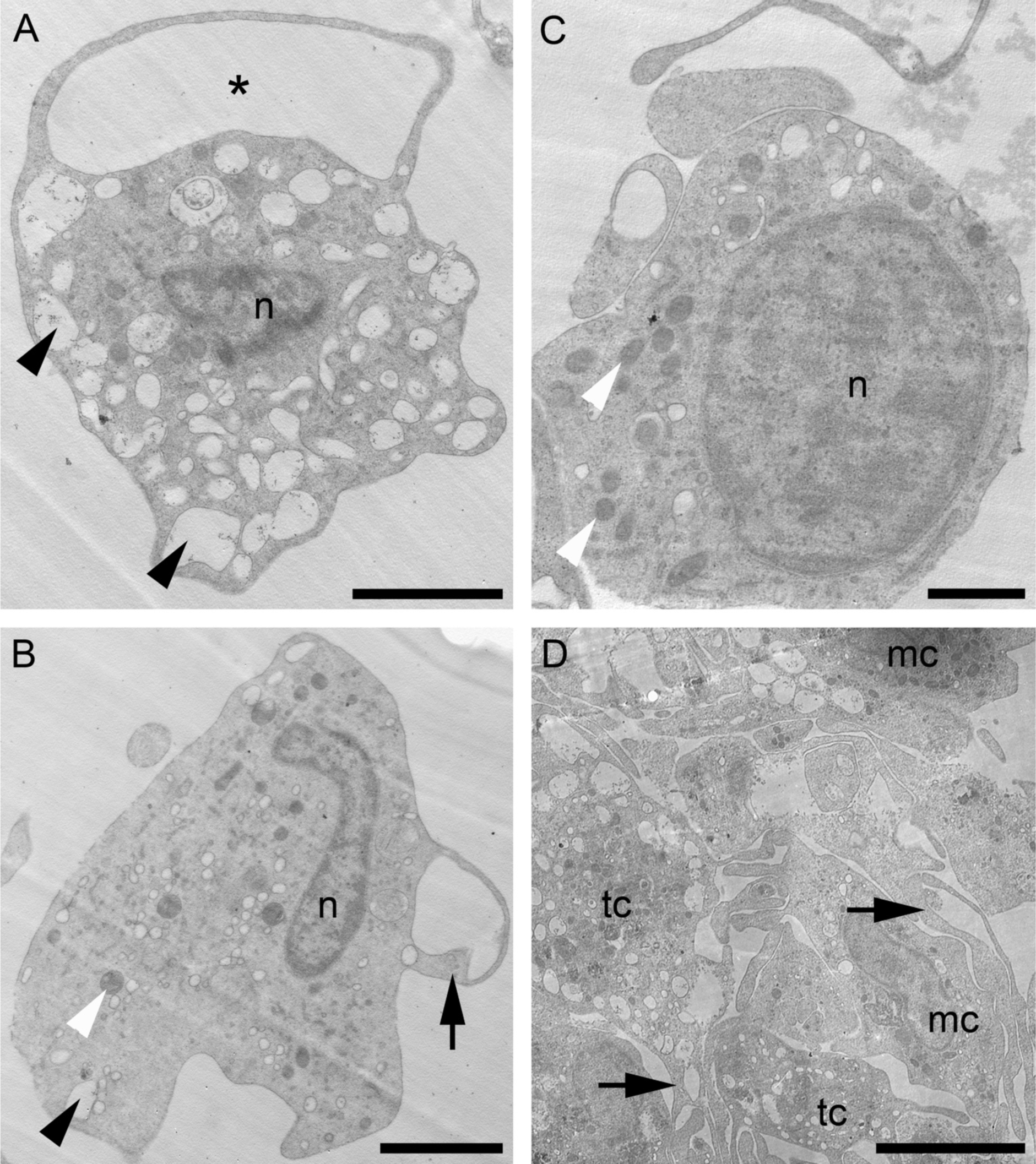
Coelomocytes morphology by transmission electron microscopy. **A**. Thrombocyte-like cell full of roundish electron-lucent vesicles (arrowheads) and with a process forming a loop (asterisk); **B**. Macrophage-like cell showing few electron-lucent vesicles (black arrowhead), mitochondria (white arrowhead) and few short cytoplasmic extensions (arrow); **C**. Slightly undifferentiated cell with a high nucleus/cytoplasm ratio and several mitochondria (white arrowheads). **D**. Aggregate of coelomocytes: both thrombocyte-like cells (tc) and macrophage-like cells (mc) possess more cytoplasmic processes packed among each other (arrows). *Abbreviations*: mc=macrophage-like cell; n=nucleus; t=thrombocyte-like cell. Scale bars: A, B=2 *µ*m; C=1 *µ*m; D=5 *µ*m.

### Characterization of cell types by fluorescence microscopy

Optical and fluorescence microscopy have been the methods of choice for the classification of echinoderm coelomocytes, mainly those based on morphological characteristics (6); (27); (33); (3). Today we understand that relevant differences exist among coelomocytes of different echinoderm classes, with some morphotypes not even represented in samples from members of the same species (3); (34). Again, disparity of protocols used for coelomic fluid collection, coelomocyte harvesting, staining conditions, sample preparation for microscopy, parameters and image processing, together with their reduced viability after collection and lack of resilience under manipulation contribute to increase the range of detected (or missed) morphological complexity. Fluorescence microscopy assays were performed using coelomocyte suspensions prepared as described for flow cytometry imaging, after adjusting the amount of dyes used in imaging flow cytometry.

Under the specific experimental conditions used for the fluorescence microscopy assays, two main morphotypes, both spherical, were identified within the smaller cell population (P1). A cell type displaying a smaller nucleus to cytoplasm ratio, where the cytoplasm is heavily granulated and surrounded by an uninterrupted bright red membrane (P1S, Figure 5A1, 2). Nuclei of the second type occupy the majority of the cell space and shows a less heterogeneous dark bright field image, completely stained blue and splashed with a reddish labelling (P1L, Figure 5A3, 4). These two morphotypes were also described in *A. amurensis* (35) and in *A. rubens* after staining with azure-eosin (27). In both studies these cells were detected in the coelomic fluid or included in the coelomic epithelium. Based on this observation, we would like to suggest that this last cell type (P1L) corresponds to a non-differentiated form of the other more mature first cell type (P1S). In line with our proposal, P1L has been suggested as the progenitor of the circulating cells (14), since they displayed no signs of differentiation and constituted 50% of the weakly attached CE cells (27). Both CE cell types showed incorporation of BrdU, although in the coelomic fluid only the cells with smaller nucleus to cytoplasm ratio presented the same BrdU-positive feature (27). P1S properties, such as proliferative activity, undifferentiated cell morphotype, ability to migrate and presence of euchromatin, are also exhibited by other animals’ stem cells (*i*.*e*. planarian neoblasts; (36)).

Using the same experimental conditions, the more abundant and bigger coelomocytes (the P2 population) were classified as phagocytes by other authors (3). The higher forward scatter determined for P2 cells (see above), compared to P1, is justified by a more complex internal cellular architecture. According to Sharlaimova and collaborators (27); (35) P2 cells incorporate BrdU suggesting their proliferative activity *in vivo*. Despite the similar cell size distribution and similar nucleus/cytoplasm diameter ratio, morphological heterogeneity has been observed within this population. We have observed diverse morphotypes with regular (Figure 5 A5), petaloid (Fig. 5 A6) or filopodial shapes (Figure 5 A7). This morphological diversity has also been described for echinoid coelomocytes (37). The regular morphotype, also referred as polygonal, presents some vesicles, mainly around the nucleus, and a spot of red staining by FM4-64 (membrane staining dye), which is frequently seen in the same cellular region (Figure 5 A5). The other two morphotypes present clearer cytoplasm and are almost not labelled with FM4-64. The petaloid form, named due to the presence of three-dimensional loops and convoluted plasma membrane foldings, undergoes a rapid conversion to the filopodial form when activated by the contact with foreign particles, through manipulation or simply as the result of contact with air (6). Chia and Xing (1996) (38) reported that the transformation from the petaloid to the filopodial forms is associated with the activation of phagocytic processes in sea cucumber *H. leucospilota*. This change occurs as a result of readjustments of the actin cytoskeleton (6); (15). Henson and collaborators (2015) (39) showed that the inhibition of arp 2/3 complex, the actin filament nucleator and branch inducer, in suspended echinoid coelomocytes drove a lamellipodial-to-filopodial shape change, and during cell spreading a new structural organization of actin was generated. Our own results are consistent with the observations of Coteur and collaborators (2002) (19), which, besides some dissimilarities in the FACS profiles, point to the presence in the coelomic fluid of a cell type with lower size and complexity (P1) plus three morphotypes of larger cells (P2; regular, petaloid and filopodial). According to Coteur and collaborators (2002) (19) and our FC results, we suggest that P1 cells are not involved in immune response, while petaloid and filopodial cells respond strongly to bacterial infection first and later on show a marked decrease during the period post-infection.

In addition to these P2 cell types, three other morphotypes, not previously described in the bibliography, were systematically observed in *M. glacialis* samples. One of them (Figure 5 B1) is roundish to oval in shape (10 μm x 15 μm), surrounded by a membrane showing irregularly distributed bright dots. The nucleus has a round shape, is located in the centre of the cell and usually has nearby dark structures surrounded by membranes coloured by FM4-64. The rest of the cell is colourless and typically granulated. Figure 5 B2 shows a representative of the second new morphotype. It presents similarities with the polygonal phagocytes, although in this case with a concentrated granular region that includes the nucleus surrounded by homogenous cytoplasm that leads to a very elongated filopodium (typically 20-30 μm) terminally subdivided into a series of very thin filopodia. Only through imaging FC we were able to detect a potentially new P2 spherical cell morphotype that shows heterogeneous cytoplasm and nucleus (Figure 4 D4). In this cell type FM4-64 stain seems to emphasize the heterogeneities of the cytoplasm.

### Coelomocyte ultrastructure

In the fluid of *M. glacialis* we have identified by TEM analyses two main cell types of free-wandering coelomocytes, that we associate to the P2 population. The first coelomocyte type (Figure 6A) presents an irregular shape, with numerous long and thin cytoplasmic processes. They have a mean diameter of about 5-7 μm. The nucleus, about 2.5 μm in diameter, has often an irregular shape and is mainly euchromatic, with the heterochromatin concentrated in scattered spots and in the periphery. They often present cell processes forming large loops which are typical of the inactive form of the petaloid sub-population of coelomocytes (38); (7); (3) (Figure 6A). The filopodial “activated” form was mainly found in coelomocyte clots/aggregates where they presented an increased number and length of cytoplasmic processes, as described by Gorshkov and collaborators (2009) (15). The cytoplasm is filled by numerous electron-lucent vesicles of different sizes: most of them appear optically empty though a few contain a fine granular material (Figure 6A). Only very occasionally they present small phagosomes in the cytoplasm. From an ultrastructural point of view, these cells are very similar to the free “mature coelomocytes” of *A. rubens*, described by Gorshkov and collaborators (2009) (15). Therefore, and according to our results, the petaloid/filopodial coelomocytes are two functional states of a same cell type which is actually specialized in clotting processes rather than participating in phagocytic activities. In vertebrates the haemostatic function is performed by highly specialized cells as thrombocytes/platelets which, once activated, undergo similar drastic shape changes, including the extension of numerous filopodial processes (40); (41), as also described for the petaloid-filopodial transition of coelomocytes. Therefore, we consider this latter the functional (and partially morphological) analogous of thrombocyte-like cells. According to Levin (2007) (42), the first cells specialized in haemostasis probably appeared in non-mammalian vertebrates. Nevertheless, our results open the intriguing possibility that this specialization have in fact occurred (or originated) at the base of the deuterostome lineage.

The second coelomocyte type also presents a heterogeneous aspect and an irregular shape (Figure 6B). It also has similar cell body and nucleus sizes and aspect, although without heterochromatin scattered spots. These cells usually bear relatively few, and quite short, cytoplasmic extensions, resembling pseudopodia. The cytoplasm is clearly less electron-dense than that described for the petaloid/filopodial cytotype and it contains numerous phagosomes, different inclusions, heterogeneous in size and appearance (small oval-shaped electron-dense inclusions and few little roundish electron-lucent vesicles), and numerous mitochondria. These cells have few large vesicles containing fine granular material, which is mainly located in the peripheral cytoplasm, and presenting also a well-developed Golgi apparatus. Overall, their ultrastructure clearly differs from that of the petaloid/filopodial cytotype previously described. However, as with the petaloid/filopodial cytotype, they undergo a morphological transition (longer pseudopodia, irregular cell and nuclear shape, and higher number of phagosomes) when activated. Considering their ultrastructural features, these cells are apparently more specialized for the phagocytic activity, therefore representing the functional analogous of the vertebrate monocytes/macrophages. The activation of these phagocytes/immunocytes is essential for the inflammatory and immune response during wound healing, as it is also known for all metazoan macrophages (43). In fact, the activated macrophages, not only phagocytize foreign material and cell debris, but also secrete a number of factors like pro-inflammatory cytokines and growth factors, such as the TGF-β (44).

Both coelomocyte types can be found free-wandering in the fluid or in aggregates (Figure 6D); when in aggregates, they are both mainly present in their respective active forms. The latter are particularly evident in the coelomic fluid of regenerating starfish (*personal observations*). The filopodial coelomocytes are generally detected at the periphery of the aggregate, whereas the macrophage-like cells in the inner part, thus further confirming the haemostatic role of the former. This is agreement with Smith *et al*., 2018 (3), who reported that phagocytes tend to aggregate in clots during wound healing, clots in which other coelomocyte types are either trapped or even contribute to their formation (30). These aggregates were also observed in our FACS experiments (Figure 1A). These heterogeneous aggregates of activated coelomocytes resemble the clot formed by these cytotypes in the wound area after 24 h following arm amputation in the same starfish species (*personal observations*). Furthermore, Ben Khadra and collaborators (2015) (9) already described a net-shaped syncytium of phagocytes covering the wound area of an amputated arm of the related starfish *E. sepositus*.

Among the free-wandering coelomocytes, we have also observed the sporadic presence of a third cytotype, possibly corresponding to the P1 population. This presents a more undifferentiated aspect with a cell body of 2-3 μm in diameter, a high nucleus/cytoplasm ratio and, occasionally, few short processes branching from it resembling those found in petaloid/filopodial cells (Figure 6C). The nucleus is mainly euchromatic, with a diameter of around 1.5 μm and few electron-lucent vesicles, again similar to those found in thrombocyte-like petaloid/filopodial cells, can be present in the cytoplasm. The cytoplasm staining and appearance of this latter cell type also resembles the clotting coelomocytes previously described. These undifferentiated cells could correspond to the undifferentiated “lymphocytes” often described in echinoderms (3)and can be the progenitors of at least this coelomocyte sub-population, as suggested for the undifferentiated cells of *A. rubens* (45) or even be stem cells with a relevant role in starfish regeneration processes.

## Conclusions

The present study constitutes the first attempt at cataloguing all cell types present in the coelomic fluid of the asteroid *M. glacialis*, native to the eastern Atlantic Ocean, using an innovative integrated strategy that combines different methodologies: Flow Cytometry, Imaging Flow Cytometry, Fluorescence Microscopy and Transmission Electron Microscopy. The results of this work suggest that *M. glacialis* coelomic fluid has two different coelomocyte populations, here named P1 and P2, which have different sizes and morphologies, and that are present at different relative abundances. The detailed study of cellular properties shows that P1 and P2 cells differ in ploidy and cell viability, as well as in in the cell cycle parameters. From light microscopy observations, the most abundant cell population, P2, has the appearance of the already described echinoderm phagocytes, including the three known morphotypes (regular, filopodial and petaloid). P1 is constituted by the smallest group of cells present in the coelomic fluid with two differentiable morphotypes; they are able to incorporate exclusion dyes, right after their extraction from the organism, indicating that they have a low *ex vivo* viability.

In the recent literature, it is stated that 95% of starfish coelomocytes present clotting and phagocytic activity, being classified as a single population called phagocytes/amoebocytes (3). From an ultrastructural point of view, our results confirm that the abundant P2 population can be actually ascribed to two distinct morpho-functional cytotypes, differentially involved in clotting phenomena or phagocytosis. These cells tend to aggregate, namely after clotting induction. Additionally, two other morphotypes were only detected by fluorescence microscopy and a third one when using a combined microscopy/FC approach, revealing the importance of combining various approaches to characterize CF cellular composition. Mainly based on our ploidy, cell cycle and morphology results, we suggest that the several P2 cytotypes described above are parts of morphologically and functionally differentiated sub-populations, possibly originating from a P1 progenitor pool. This hypothesis is in accordance with Gorshkov and collaborators (2009) (15) who also proposed that some cell morphotypes could correspond to diverse differentiation or activation stages.

Our work here points to the need of a further systematic characterization of the histology/physiology of these cell morphotypes, which should be instrumental in ascertaining the origin and lineage relationships of those cells.

Before ending this manuscript, it is important to point out that this project allowed us to implement optimized experimental protocols to study different coelomocyte characteristics, such as cell viability, cell cycle or cell morphologies with the aid of the powerful imaging FC technologies. We envision that this experimental approach will be especially useful in the future elucidation of coelomocyte functional competences. Moreover, the combination of the above technologies with transcriptomics and proteomics studies suggests a way to unravel and/or confirm some of their putative role, also providing us with a powerful tool to discover specific cellular biomarkers.

## Supporting information

Figure S1 Flow cytometric analysis of circulating coelomocytes using the anticoagulant buffer

## Acknowledgments

The authors acknowledge Vasco da Gama Aquarium (Dafundo, Oeiras, Portugal), namely Dr. Fátima Gil and Miguel Cadete, for starfish maintenance. To Maristem COST Action (CA16203) supported by COST (European Cooperation in Science and Technology) for funding PM STSM (February 2019) at AVC Laboratory. Electron microscopy assays were carried out at NOLIMITS, an advanced imaging facility established by University of Milan.

## Author’s contributions

AVC, MS, RG and RZ developed the experimental design. BO, CB, CA conducted the flow cytometry experiments, supervised by RG. BO, JR and AVC were involved in flow cytometry imaging experimental work. SG, CF and MS performed the electron microscopy assays. BS and AVC performed the fluorescence microscopy experiments. All authors contributed to the analysis of the experimental results and to the manuscript. The whole project was coordinated by AVC.

