## Supplementary material for "A combined characterization of coelomic fluid cell types in the spiny starfish *Marthasterias glacialis* – inputs from flow cytometry and imaging": Figure S1 Flow cytometric analysis of circulating coelomocytes using the anticoagulant buffer

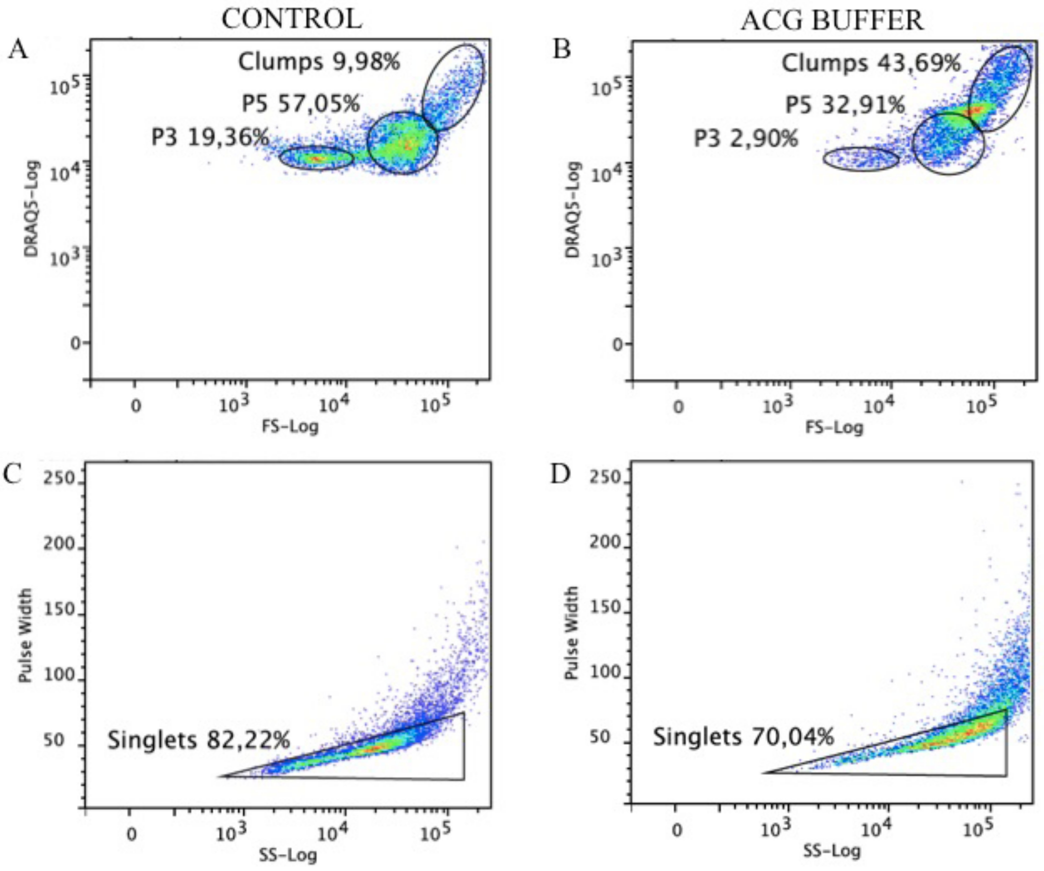


**Figure S1 Flow cytometric analysis of circulating coelomocytes using the anticoagulant buffer** (0.5 M NaCl, 5 mM $MgCl_2$, 20 mM HEPES and 1 mM EGTA pH=7.5)**.** Coelomocytes were stained with DRAQ5**. A, C.** Dot plots representation of **(A)** coelomocyte populations and **(C)** its singlets gated without using anticoagulant solution. **B, D.** Dot plots representation of anticoagulant solution effect in **(B)** coelomocyte populations and **(D)** its singlets gated
